# Drivers of host-pathogen community assemblies in European forests and urban green spaces

**DOI:** 10.1101/2024.09.19.613856

**Authors:** Vincent Sluydts, Marie Bouilloud, Maxime Galan, Hussein Alburkat, Anaïs Bordes, Vincent Bourret, Valeria Colombo, Luc DeBruyn, Lara Dutra, Jana Eccard, Jasmin Firozpoor, Romain Gallet, Maciej Grzybek, Heikki Henttonen, Jens Jacob, Andrew McManus, Tarja Sironen, Peter Stuart, Caroline Tatard, Benjamin Roche, Herwig Leirs, Nathalie Charbonnel

## Abstract

Major advances in the understanding of infectious diseases have been achieved in the last decades. However, the persistence and re-emergence of pathogens continue to raise public and veterinary health concerns, of which the recent COVID-19 pandemic may be one of the most dramatic examples. Understanding the impact of habitat alterations and concomitant biodiversity loss on pathogen transmission and emergence from wildlife remains challenging. Here, we aim to elucidate the interlinkages between biodiversity and rodent-borne diseases at local and European scales. We present recently collected host-pathogen data from 21 temperate forest sites and eight urban green spaces throughout five European countries, environments where rodents are abundant and human/domestic animals – wildlife interactions are likely to occur. 3766 specimens were analyzed during the period from 2020 to 2022 comprising 15 different small mammal species. Different organ tissues of each specimen were screened for bacteria by either 16S rRNA amplicon sequencing or specific PCR. The presence of antibodies to different families of viruses was screened using immunofluorescent assays. A multitude of pathogens of zoonotic potential from several genera including *Bartonella, Borrelia, Mycoplasma, Anaplasma, Neoehrlichia, Leptospira*, Orthohantavirus and Orthopoxvirus were detected at non-negligible prevalence in 11 different terrestrial mammal species. A shift in host community composition was observed along the anthropization gradient with more urban adapters in more anthropized sites. Pathogen richness increased with an increase in host species diversity, following the “host-diversity begets parasite-diversity” hypothesis. The absence of some vector-transmitted parasites in urban areas suggests a shift in pathogen community along the anthropization gradient. Host species and host intrinsic factors were dominant explanatory variables for endoparasitic *Mycoplasma* species and *Sarcocystidae*, while extrinsic environmental and climatic factors where influential in explaining variations in occurrences of several vector-transmitted pathogens. *Apodemus sylvaticus* and *Clethrionomys glareolus* were important connector host species in respectively urban green spaces and temperate forests. Increased host diversity, but not anthropization, correlated with a richer pathogen community. These results ultimately lead to an increased understanding of the complex host-pathogen system at the local landscape that can aid future management decisions and support the public health sector.

## Introduction

The persistence and re-emergence of pathogens pose significant public and veterinary health concerns worldwide. The recent COVID-19 pandemic serves as a stark reminder of the complex interplay between wildlife hosts and their zoonotic agents [1]. Consequently, understanding the impact of habitat alterations and biodiversity loss on pathogen diversity, transmission, and emergence from wildlife has become an urgent research priority [2].

Local assemblies of host-pathogen communities are the culmination of complex processes operating at different spatial [3] and temporal [4] scales. Coarse scale processes like speciation, species sorting, and environmental filtering interoperate with higher resolution processes such as biotic interactions, host and vector dispersal, pathogen transmission, stochasticity and individual immunity [5,6]. Together these processes define a regional host-pathogen pool in a local landscape at a given time. A key question in disease ecology is to what extent these processes drive the geographical distribution of host-pathogen interactions and contribute to assembly patterns of pathogen communities at the local landscape level.

From a pathogen’s perspective, a given host functions as a mobile resource patch [7]. High host diversity leads to a rich and variable host community, leaving ample opportunities for pathogens to colonize these new habitats. This concept is called the “host-diversity begets parasite-diversity” hypothesis and predicts pathogen diversity to increase with host diversity [8]. On the other hand, the dilution effect hypothesis suggests that a high level of biodiversity tends to “dilute” competent hosts within host community, thus limiting pathogen transmission [9]. Under this scenario, and according to the fact that competent hosts tend to be those that remain or colonize following biodiversity loss, it is expected that such loss would increase the average host community competence.

The interaction of aforementioned mechanisms and processes shapes local pathogen-host assemblies [10]. The complexity of multi-host, multi-pathogen ecosystems and the lack of empirical data across different gradients at a local landscape scale have inhibited our predictive understanding of the system [3]. To comprehend how environmental shifts impact host-pathogen assemblages, it is imperative to adopt a holistic, multi-species perspective [3].

Antropization is one of the major global factors threatening wildlife worldwide. In this study, we define anthropization or human influence as the human footprint index [11]. An increase in anthropization is associated with habitat fragmentation and increased pollution, with strong impacts on biodiversity, including abrupt shifts in community composition. Under those conditions, synanthropic host species, generally known for their host competence, tend to thrive [12], increasing host community competence and risk of infection [9]. As a result, human influence changes patterns of disease transmission and emergence by shifting host and vector community composition, thereby reshaping the competence, assembly, and interactions of host-pathogen communities [9,13].

Several meta-analyses have emphasized the role of anthropization in zoonotic emergence. Jones et al. (2008) [14] demonstrated that disease emergence is largely mediated by anthropogenic changes, with Gibb et al. (2020) [15] and Murray et al. (2019) [16] finding that human-influenced areas and urban wildlife host more diverse and abundant zoonotic pathogens. Albery et al. (2022) [17] attributed this pattern to greater overall pathogen diversity in urban settings rather than an increase in zoonotic pathogen richness. To further disentangle the role anthropization plays in increasing the risk of zoonotic infections through altering the multi-host, and multi-pathogen assemblies at the local landscape scale, empirical studies are needed [17].

This present study focused on rodent-borne zoonotic pathogens *sensu lato* [18], due to their significant implications for public health and veterinary medicine [12]. Moreover, the (re-)emergence of several major rodent-borne zoonotic diseases seems to be associated with urbanization (e.g. [19,20]). Therefore, we sampled small mammal host communities in urban green spaces and forest fragments along an anthropogenic gradient throughout five different countries across Europe.

We aimed to describe the metacommunity diversity and structure [10] of rodent-borne pathogen communities from these urban green spaces and temperate forests in Europe. We hypothesized that our semi-experimental filter of anthropization leads to a reduction in host species diversity, with an increase in those hosts adapted to a more urban environment (referred to as “urban adapters” herein), generally having a faster pace-of-life phenotype [9] and acting as more competent reservoirs [15]. These changes in host community competence are thought to affect the pathogen community in predictable, albeit opposing ways. We hypothesized that anthropization leads to an overall loss in host species diversity, which covaries with overall pathogen richness, due to the mechanisms of resource availability and/or habitat heterogeneity [2]. On the other hand, we hypothesized that anthropization gives rise to a turnover of host species and that synanthropic competent hosts become more prominent, leading to an increased diversity and prevalence of pathogens.

The observed host-pathogen community was studied in three ways. First, we compared the host community structure per habitat and described specific infection and co-exposure patterns, checking for non-random co-occurrence patterns between pairs of infection. Second we used a network analysis to investigate the difference in host-pathogen network structure between temperate forest fragments and urban green spaces and to assess the relative epidemiological importance of rodent host species [21]. Modifications in the network structure resulting from habitat alteration can help assess changes in pathogen transmission providing a better understanding of the drivers behind zoonotic hazards [22]. We expected less modular but more connected networks due to fragmentation and generalization, and a high centrality for urban adapters in the urban green spaces [23]. In addition, we expected to observe an increase in nestedness along the anthropization gradient resulting from an increase in habitat fragmentation and lower availability of favourable hosts [24].

Finally, we investigated biotic and abiotic filtering processes shaping local assemblies and individual hosts’ fitness associations with the pathogen community. Using a joint species distribution framework, we examined how these processes influence pathogen community composition, accounting for transmission traits and spatiotemporal variation. We considered environmental, climatic, and anthropogenic drivers as abiotic factors and individual host characteristics and host community diversity as biotic factors. This comprehensive analysis describes the relative importance of various factors in pathogen community assembly and remaining pathogen associations, enhancing our understanding of the complex host-pathogen system at the local landscape level to inform management decisions and support public health efforts.

## Material and Methods

### Trapping sites and host data

Data were collected during Spring and Autumn 2020-2022 in five countries in two habitats: temperate forests and urban green spaces [Figure 1]. Host specimens were captured using live and snap traps with the aim of collecting up to 25 specimens per species in each of three sites per habitat per country. The trapping protocol is described in detail for France by Pradel et al., (2022) [25]. It is adapted to *in-situ* ethical standards and to allow for the use of local traps and baits. In short, a variety of baits and traps: INRA [France], Sherman live [Belgium], Longworth [Ireland & Germany] and Ugglan [Germany] were used to sample small mammal communities. In addition, rat-live traps [Belgium and France] and snap traps [Germany] were used.

**Figure 1:**
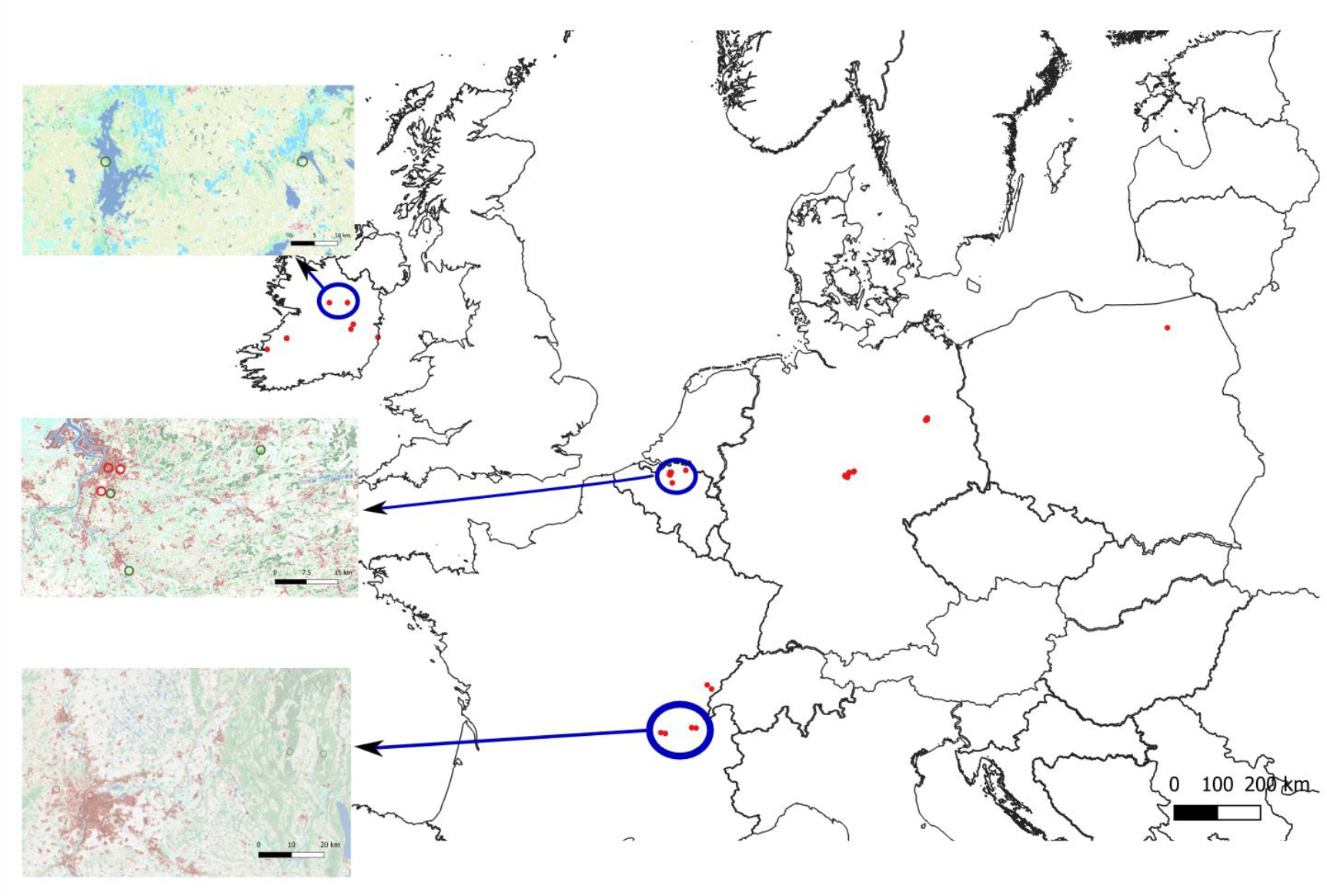
Overview of trapping sites [red dots] in Europe. Insets show examples of sites in Ireland, Belgium and France with Copernicus layers for imperviousness, water, forest and grasslands, and urban green spaces as red circles and temperate forests sites in green.

Trapped specimens were euthanized and dissected immediately after capture. Morphological features, including body length and weight, sex, and sexual maturity, were recorded. Species identification was based on morphometrics and confirmed by molecular analyses when necessary. Several organ samples were taken for pathogen detection and gut microbiome characterization incl. blood, spleen, kidney, and colon [25].

Animal capture and handling have been conducted according to the local and European regulations on care and protection of laboratory animals, a more detailed description can be found in the data papers [25,26].

### Pathogen data

#### Detection of Orthohantaviruses, Orthopoxviruses and Mammarenaviruses

Exposure to Orthohantavirus, Orthopoxvirus and Mammarenavirus was detected using direct immunofluorescence assays (IFA) on blood serum from trapped specimen. The assay detects IgG antibodies, indicating long-term immune response. Assays were performed as described by Kallio-Kokko et al. (2006) [27] using slides coated with Vero E6 cells infected with i) Puumala (PUUV) or Dobrava virus (DOBV) for Orthohantaviruses, ii) cowpox virus to detect Orthopoxviruses, and iii) lymphocytic choriomeningitis virus (LCMV) for Mammarenaviruses.

#### Pathogenic Leptospira spp. detection

Genomic DNA was extracted from specimens’ kidneys using 96-well plate animal genomic DNA extraction miniprep kits (Biobasics©). DNA was eluted in 150 μl. Pathogenic *Leptospira* spp. detection used real-time polymerase chain reaction (qPCR) targeting the lipL32 gene on a LightCycler® 480 (Roche Diagnostics, France), following Dobigny et al. (2015) [28]. Rodent kidney qPCR was performed in duplicate using 96 or 384-well microtiter plates, with 2 μl DNA in 10 μl final volume per reaction. Plate included negative controls (for extraction and qPCR) and positive controls. Absence of amplification in at least one duplicate indicated absence of leptospirosis infection.

#### Pathogen bacterial community detection analyses

Bacterial DNA was extracted from spleen samples using the DNeasy Blood & Tissue kit (Qiagen). We amplified and sequenced a 251-bp fragment of the 16S rRNA V4 region using a modified version of the dual-index method [29], as detailed in Galan et al. (2016) [30]. Each extraction was analyzed in duplicate across 20 MiSeq runs.

Sequences were processed using FROGS pipeline [31] to generate an OTU (Operational Taxonomic Unit) abundance table. Taxonomic affiliation was obtained using the Silva database v138.1 with RDP Classifier [32] or blastn+ [33]. False positives were filtered as per Galan et al. (2016) [30]. Only OTUs confirmed in both replicates were retained. Following Abbate et al., (2024) [34], analysis included only OTUs with ≥ 500 reads across all samples, and established pathogenicity from the literature.

### Biotic and abiotic covariates

#### Characterization of host gut microbiome

Host gut bacteriota from colon samples were characterized using 16S barcoding. The dada2 pipeline (Qiime2_2021.11) identified amplicon sequence variants (ASVs). False positives were filtered per Galan et al. (2016) [30]. Alpha diversity of each specimen’s gut bacteriota was calculated using specific richness (log transformed), as this is expected to correlate with fitness and to decrease in disturbed environments [35,36].

#### Characterization of site environmental, climatic and anthropization variables

Environmental data were extracted from the Copernicus land monitoring service [https://land.copernicus.eu/] as percentage of land-use class in a 1 km radius around each sampling site. Percent of broad-leaved and coniferous forest, grassland, imperviousness, and water, both permanent and temporary, was extracted (Figure 1 - insets).

Climatic variables were extracted from daily gridded meteorological data for Europe [https://cds.climate.copernicus.eu/]. Maximum temperatures the previous summer and minimum temperatures the previous winter within a 1 km radius were extracted [37]. Total rainfall in the previous year was accumulated over the same buffer range for each site.

Anthropization is quantified using the Human Footprint Index (HFI), a composite measure that combines population density, infrastructure, accessibility, and energy deployment, available at a 300-meter resolution globally as provided by the Wildlife Conservation Society [38]. The degree of anthropization was determined by calculating the average HFI within a 1-kilometer radius around each site.

#### Characterization of host species biodiversity

The diversity of host communities was extracted from small mammal biodiversity models for Europe provided by Wint et al. (2013) [39]. We opted for the model variant which included 10 different small mammal species and the variable extracted was named Host Species Diversity [HSD]. As the resolution of these models is coarser compared to earlier variables, we decided to use the minimal non-zero value from a 10 km radius as a metric.

### Statistical analyses

All statistical analyses were performed in R v4.3.1

#### Host community and anthropization

We analyzed host community composition changes with anthropization by calculating β diversity using Cao dissimilarity between sites [40]. Differences between habitats and countries were tested using PERMANOVA with the adonis2 function (R package vegan [41], 1000 permutations). Host specimens were categorized as urban “avoider”, “adapter” and “dweller”. Life history traits were extracted from Plourde et al. (2017) [42] and Albery et al. (2022) [17], resulting in two mass-corrected principal components explaining 86% of the variation in six mammalian traits. The first component corresponds to a general fast-slow continuum, the second is more oriented towards gestation time and larger offspring [17,42]. An ANOVA tested differences in these traits between different wildlife responses to anthropization.

#### Infection and co-infection/co-exposure patterns

“Co-infection” was defined as the concomitant infection of a host with bacteria detected directly by 16S metabarcoding or conventional PCR techniques. “Co-exposure” is used when a host is or was infected also by a virus detected by serology (IFA). To determine if pairs of co-infection/co-exposure occurred more or less frequently than random, the cooccur package in R was used. In essence a probabilistic model was applied which calculates expected frequencies of co-occurrences under assumption of a random distribution. After comparison with observed frequencies, we could determine if pairs of co-occurrences appeared more or less frequently in these data [43].

#### Host-pathogen network statistics

All networks were constructed with the bipartite package [44] to examine (1) the relationship between host communities and anthropization, (2) the composition of hosts and pathogens communities in different habitats. To assess differences in metacommunity structure between urban green spaces and temperate forests, a host-pathogen network was built for each of these habitats. We hypothesized that anthropization should lead to a decrease in host community diversity and an increase in species adapted to anthropization, these being better reservoirs than urban avoider species. As a result, we predicted an increased infection prevalence (and hence more co-infections) as anthropization increases [19]. For each network, the bipartite package in R was used to calculate network-level statistics (connectance [C], network specialization [H2], modularity [Q] and weighted nestedness [wNODF]) [45,46]. To test the hypothesis that these measures differ between the networks, a bootstrap method was implemented in which 5,000 samples of the difference for each statistic between the two habitats were drawn based on the r2dtable null model [47]. The observed difference of each statistic was evaluated against this distribution to obtain a p-value. To evaluate which host species are important “connectors” in each network, we calculated two species-level centrality statistics: the normalized degree [ND] and the weighted betweenness [BC]. The first measures the generalization of a host species by the number of host-pathogen interactions, while the latter describes the importance of a host as a connector between different parts of the network [48].

#### Pathogen community structure and pathogen-pathogen associations

A joint species distribution model [JSDM] was fitted to the pathogen community data using the Hmsc package to infer how pathogens respond to biotic and abiotic signals, meanwhile accounting for co-occurrence patterns related to unmeasured variables [49]. In essence, the JSDM is a Bayesian multivariate generalized linear latent variable model able to fit pathogen communities (as opposed to single-species models) and accounting for fixed and random covariates as well as trait and phylogenetic effects [50]. As a response variable, the pathogen presence-absence data detected by either serology, qPCR or 16S barcoding approaches, for the host genera *Apodemus* and *Clethrionomys* as they occurred at all trap sites except Poland, were modelled using a probit regression. Only pathogens with at least 50 occurrences were kept in the analysis. As explanatory covariates we opted for intrinsic, extrinsic, and anthropogenic covariates. As intrinsic variables we considered host individual characteristics (sex, weight, sexual maturity, gut microbiome diversity) and host species. As extrinsic covariates, both site-level environmental (percent of grassland, broadleaved and coniferous forest, and temporary and permanent water bodies) and site-level climatic (minimum temperature past winter, maximum temperature past summer and total accumulated rainfall over the past year) covariates were considered. Anthropization was modelled by means of the human footprint index [HFI] and host species diversity [HSD] [39]. The specific transmission route of each pathogen [i.e. vector, environmental or direct] was considered in the trait matrix. Different JSDM variants [Table 1] were fitted to the data.

**Table 1:** Overview of joint species distribution models (JSDM) fitted to the data. Sex, sexual maturity, weight, log-transformed gut microbiome richness (collectively referred to as [HOST]), and species were the host intrinsic characteristics. Climate [CLIM] and environment [ENV] were considered as extrinsic covariates. Anthropogenic effects were host species diversity [HSD] and human footprint index [HFI]. Spatial [SP] and temporal [TMP] level random effects were considered. Mode of transmission, categorized into direct, vector or environment, was accounted for as trait effect. Explanatory performance was evaluated based on mean Tjur R^2^. Predictive performance was evaluated on three-fold cross-validated mean Tjur R^2^.

| MODEL | Fixed Effects | Random Effects | Trait | Explanatory performance | Predictive Performance | WAIC |
| --- | --- | --- | --- | --- | --- | --- |
| HAB | HABITAT | SP + TMP | TRANS | 0.132 | 0.126 | 3.18 |
| INT | HOST + SPECIES | SP + TMP | TRANS | 0.212 | 0.194 | 2.82 |
| EXT | CLIM + ENV | SP + TMP | TRANS | 0.143 | 0.130 | 3.16 |
| ANT | HSD + HFI + HSD:HFI | SP + TMP | TRANS | 0.136 | 0.125 | 3.17 |
| INT+EXT+ANT | HOST + SPECIES + CLIM + ENV + HSD + HFI | SP + TMP | TRANS | 0.224 | 0.204 | 2.79 |

We assessed the impact of intrinsic, extrinsic, and anthropogenic variables on pathogen communities, accounting for spatial and temporal variations and pathogen transmission traits. Our baseline model [JSDM M0, Table 1] compared urban green parks to temperate forests. We then evaluated models incorporating various factor combinations [Table 1]. Model selection was performed by means of WAIC, explanatory and predictive performance based on Tjur R^2^ and three-fold cross-validation [51]. We employed Hmsc v 3.0-13 with default priors [52], using four MCMC chains to generate 20,000 posterior samples. Convergence was verified through potential scale reduction factor and visual chain inspection.

## Results

### Host community and pathogen detection

During the study period, 3766 small mammal specimens were collected, identified, and analyzed (Figure 2), comprising 15 species of which 12 were rodent species [*Muridae, Critetidae, Gliridae* and *Sciuridae*] and three were shrews [*Soricidae*]. A complete set of individual host characteristics, gut microbiome and pathogen data was obtained from 3463 [92%] individuals. Rodent-specific biodiversity per field site, ranged from 1 species (Ireland) to 7 (Poland, Germany), whereas the occurrence of the different pathogen genera ranged from 5 (Ireland) to 16 (Germany).

**Figure 2:**
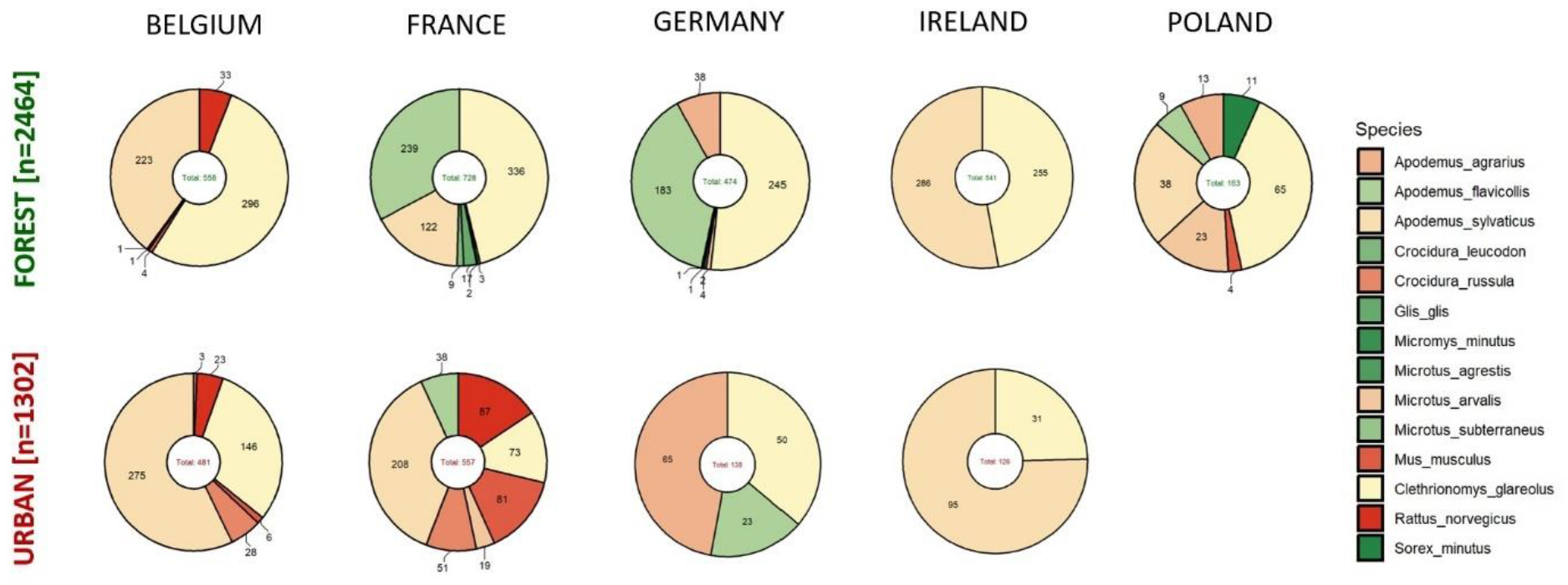
Overview of the different host species included in the host-pathogen data analyses from each country in temperate forests and urban green spaces, with indication of the number of samples collected for each specimen. Colour codes range from green, over yellow to red, following the ecological gradient from urban avoider [green], through urban adapter [yellow] to urban dweller [red].

Among the captured rodent species, the bank vole (*Clethrionomys glareolus*) and the wood mouse (*Apodemus sylvaticus*) were encountered most often, occurring in both urban green spaces and forested habitats. These species can be regarded as urban adapters [53], with *Apodemus sylvaticus* being more explorative, occurring even in the center of Antwerp. In Germany, only one *Apodemus sylvaticus* specimen in a forested habitat was recorded, but *Apodemus agrarius* was found filling a similar niche in both urban green spaces and temperate forest habitats. *Rattus norvegicus*, being an urban dweller, was also detected in forested sites in Belgium, but those sites are embedded in a dense matrix of man-made structures. *Mus musculus*, being an urban dweller, also occurred in the temperate forest in Poland (Figure 2). In France the host species composition shifted from more urban avoiders in the forested areas to urban adapters and dwellers in the urban green spaces, but the overall observed host species richness remained similar in both habitats (Figure 2).

Overall, host community composition did not differ between habitats (F= 1.62, df = 1, p = 0.20), but differed between countries (F= 12.4, df = 3, p <0.01). An interaction term modelling that habitat differences can change with country was marginally significant (F=1.88, df=3, p = 0.073), and, as is apparent from the figure, was driven by the differences in host communities in France. Regarding life-history traits of host species, we did not find differences between urban avoider, adapter and dweller species for either a general fast-slow life-history continuum [p = 0.26, LRT = 2.71, df=2] or one focused on longer gestation times and larger offspring [p = 0.32, LRT = 2.26, df=2].

A diversity of pathogens was observed in the host community [Figure 3], 54% [n=2103] of the specimens were infected with a pathogen belonging to a genus with zoonotic potential, whereas 44% [n=1693] were infected with endoparasitic *Mycoplasma* spp. which are not zoonotic but can be pathogenic to the rodent host. *Bartonella* spp. [47%] was frequently occurring followed by Orthopoxvirus [15.8%]. Other infections with zoonotic potential occurring at a prevalence higher than 1% were protozoans from the family *Sarcocystidae* [10.2%], *Candidatus Neoehrlichia* [9.1%], Orthohantavirus [4.9%], *Borrelia* spp. [3.5%], *Leptospira* spp. [3.3%], *Anaplasma* spp. [2%] and *Francisella* spp. [1.8%]. *Orientia* spp., *Chlamydia* spp., *Streptobacillus* spp. and *Rickettsia* spp. were picked up but only on rare occasions [Supplemental Table S1]. Note that on the continental shelf island, Ireland, still 41% of all pathogens that occurred in the data were detected, while only two host species were recorded in this study (see *Study limitations*).

**Figure 3:**
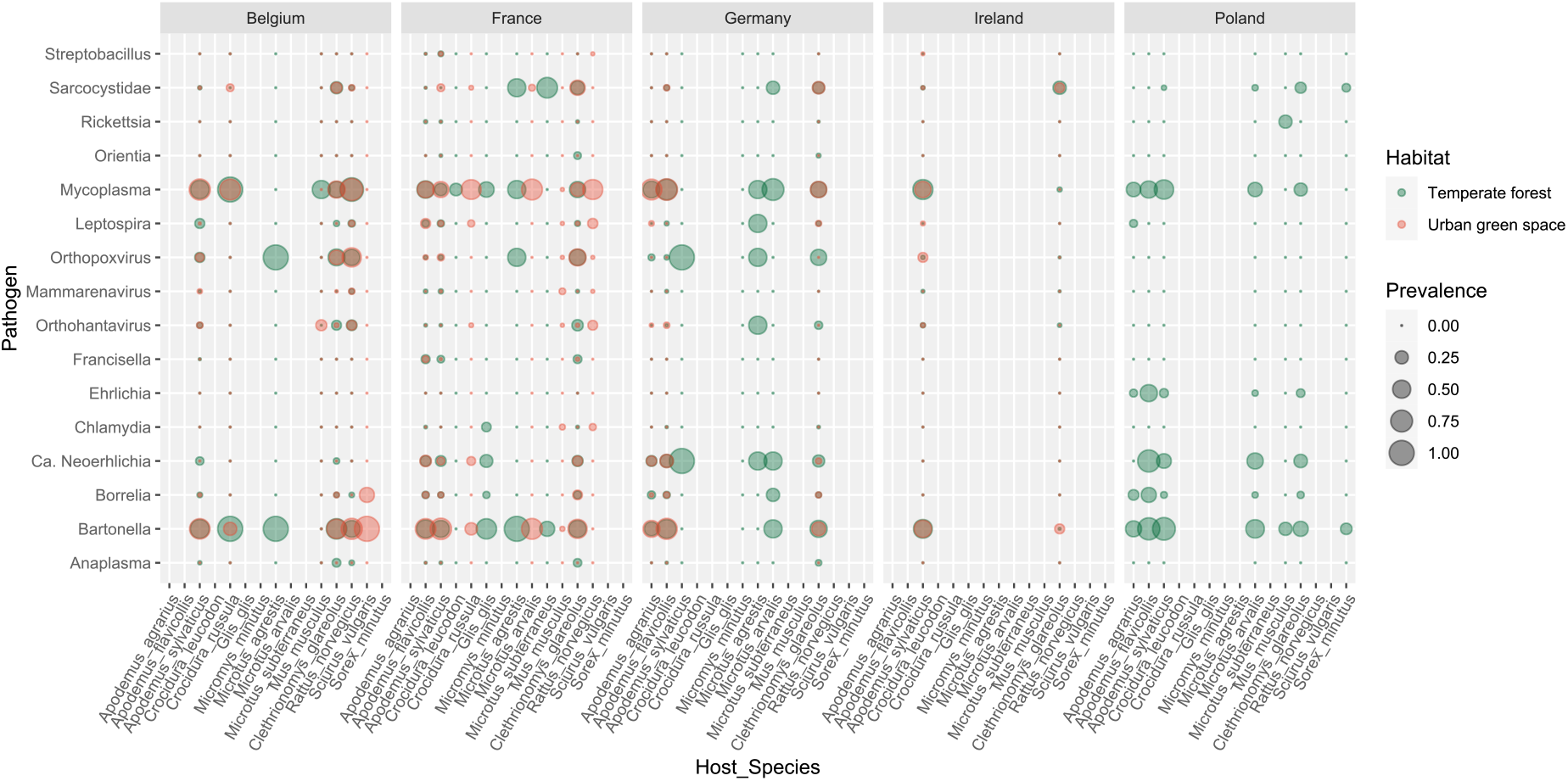
Observed pathogen prevalence for each host species in five different European countries. Green circles scale with prevalence in temperate forests, red circles with urban green spaces.

Among the different host reservoirs, high variation was detected in the prevalence of zoonotic infections and parasites that they carry or were exposed to [Figure 3]. Differences in prevalent pathogens between focal host species were observed. From the host specimens collected in lower numbers, all *Microtus agrestis* [n=5], *Sciurus vulgaris* [n=3], *Microtus subterraneus* [n=3] as well as 82% of *Glis glis* [n=17] and 27% of *Sorex minutus* [n=11] carried at least one zoonotic infection.

For the more prominently present species at least one potentially zoonotic agent was detected in 76% of *M. arvalis* [n=46], 71% of *C. glareolus* [n=1534], 68% of *A. sylvaticus* [n=1296], 67% of *A. flavicollis* [n=492], 54% of *R. norvegicus* [n=152], 47% of *A. agrarius* [n=116], 35% of *C. russula* [n=80], 15% of *M. musculus* [n=95].

### Pathogen co-infection and co-exposure patterns

Co-infection and co-exposure with at least two bacteria, protozoans or viruses with pathogenic potential [excl. *Mycoplasma*], was reported in 39% of all *C. glareolus*, 27% of *A. flavicollis*, 18% of *R. norvegicus*, 15% of *A. sylvaticus* and *M. agrestis* and 12% of *A. agrarius* [Figure 4]. Co-exposure between *Bartonella* spp. and Orthopoxvirus were predominant in *C. glareolus* and *R. norvegicus*, whereas co-occurrence between *Bartonella* spp. and *Ca. Neoehrlichia* as well as other vector-transmitted pathogens such as *Francisella* spp. and *Borrelia* spp. prevailed in *A. sylvaticus*. Significant non-random positive co-occurrences between pathogen pairs were observed in *C. glareolus* [18 pairs], *R. norvegicus* [3 pairs], *A. flavicollis* [3 pairs] and *A. sylvaticus* [2 pairs] Significant non-negative co-occurrences were found in the same host species but were less prevalent [Supplemental Figure S1].

**Figure 4:**
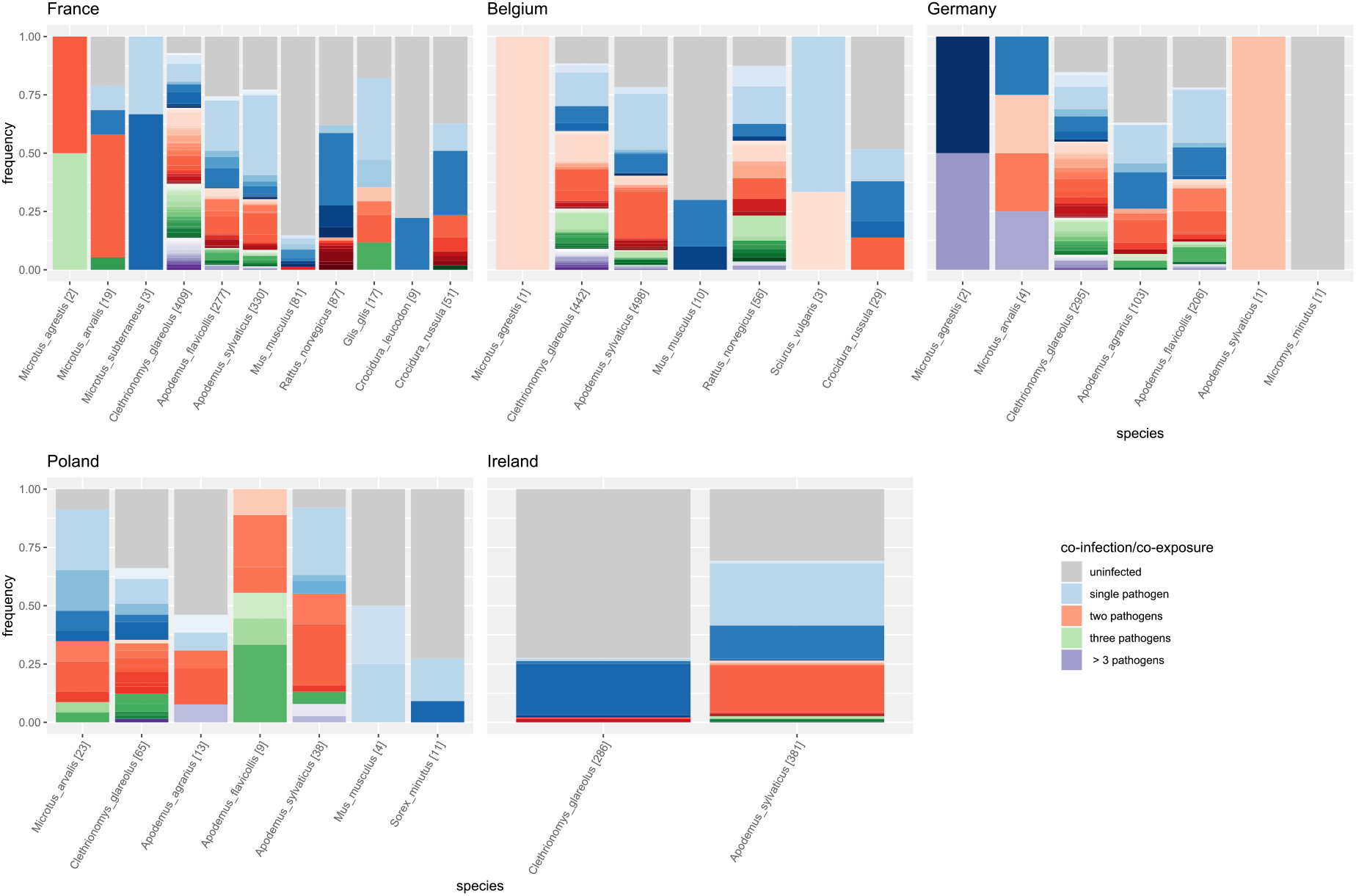
Infection, co-infection, and co-exposure patterns per host and country, proportional to the number of individuals per host [in brackets]. Grey bars indicate uninfected specimens, blue shades represent one infection, red two, green three and purple >3 infections according to the legend. Colored shades indicate different pathogens or combinations thereof.

Between country patterns of pathogen assemblies for the most common hosts, *C. glareolus* and *A. sylvaticus*, were consistent between countries, except for co-infection/co-exposure patterns in Ireland, where *C. glareolus* is invasive.

### Hosts-pathogens network

Pooling the host-pathogen interactions per habitat type demonstrated structural differences between the urban green spaces and temperate forests (Figure 5). On average, urban green spaces exhibited a more generalized network [H2, p < 0.001] with higher connectance [C, p < 0.0001], while no significant difference was observed for weighted nestedness [wNODF, p = 0.28]. Four compartments were found in each habitat consisting of different host-pathogen interactions. In urban green spaces, the largest module consisted of urban dweller species in combination with host specific *Mycoplasma* and directly or environmentally transmitted pathogens with a strong human affiliation (*Leptospira* spp., *Orthohantavirus* spp., *Mammarenavirus* spp.). Notably some vector-transmitted pathogens such as *Ehrlichia, Anaplasma, Orientia* and *Rickettsia* species appeared absent from the urban host-pathogen network.

**Figure 5:**
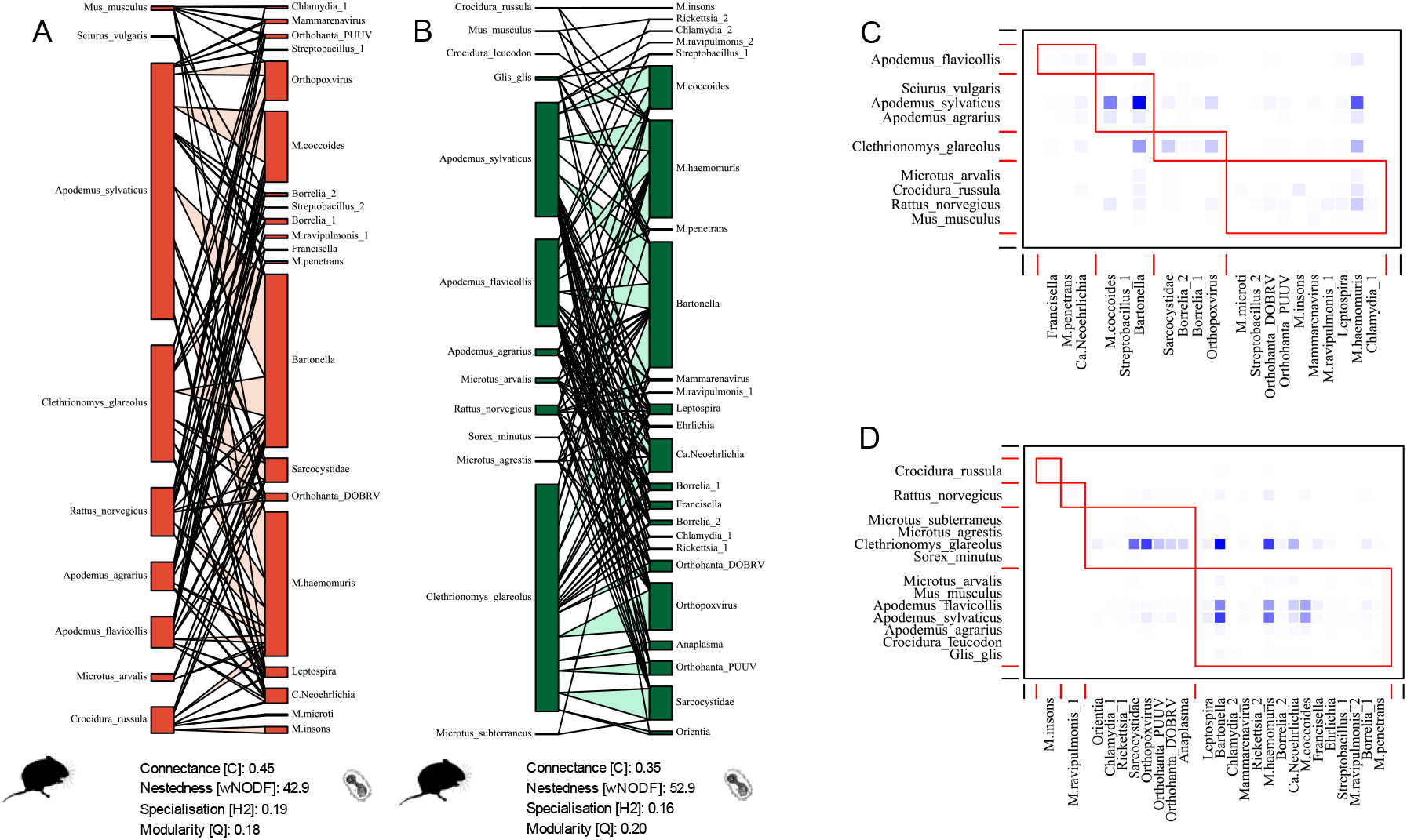
A, B: Bipartite graphs for the host-pathogen network in urban green spaces [A, red] and temperate forests [B, green] with an indication of network level statistics and C, D: modules for the urban green space network [C] and temperate forest [D] (see module description in main text). Silhouettes are taken from phylopic.org.

Ranking host species using their links with pathogen genera [ND, normalized degree] and betweenness centrality in the network [BC, weighted betweenness] revealed considerable differences between the habitats. In the temperate forests *A. flavicollis* (ND = 0.8), *A. sylvaticus* (ND = 0.8) and *C. glareolus* (ND = 0.76) showed the highest levels of generalization followed by *Rattus norvegicus* (ND = 0.48) and *A. agrarius* (ND = 0.36). However, the betweenness centrality (BC), pointed to *C. glareolus* (BC = 0.86) and *A. flavicollis* (BC = 0.14) as key central host species in the forests. In the urban green spaces, *A. sylvaticus* (ND = 0.8), *Rattus norvegicus* (ND = 0.6), *C. glareolus* (ND = 0.6) and *A. flavicollis* (ND = 0.6), followed by *C. russula* (ND = 0.4) and *A. agrarius* (ND = 0.4) possessed a high degree of generalization. Instead, *A. sylvaticus* (BC = 0.79) and *R. norvegicus* (BC = 0.21) played a key role as connectors in urban green spaces.

### Pathogen community structure and pathogen-pathogen associations

Five different models were fitted to the data to explore the contribution of intrinsic, extrinsic, and anthropogenic drivers (Table 1). The habitat baseline model [HAB] was outperformed by all combinations of intrinsic, extrinsic, and anthropogenic explanatory factors in terms of WAIC and predictive performance. Models including intrinsic host-specific characteristics provided a better fit to the data. A model combining both intrinsic and extrinsic variables with anthropogenic variables showed better explanatory and predictive power in combination with the lowest WAIC. As such it was used to show the proportion of variance explained by the fixed and random effects [Figure 6: I], and to evaluate the effect of covariates [Figure 6: II]. This model had a mean effective sample size for the fixed effects of 12,386, with an average potential scale reduction factor (psrf) of 1.007 [sd = 0.017]. For the trait effects the average psrf was 1.001 [sd = 0.0011] and for the random effects the mean was 1.012 [sd = 0.011].

**Figure 6:**
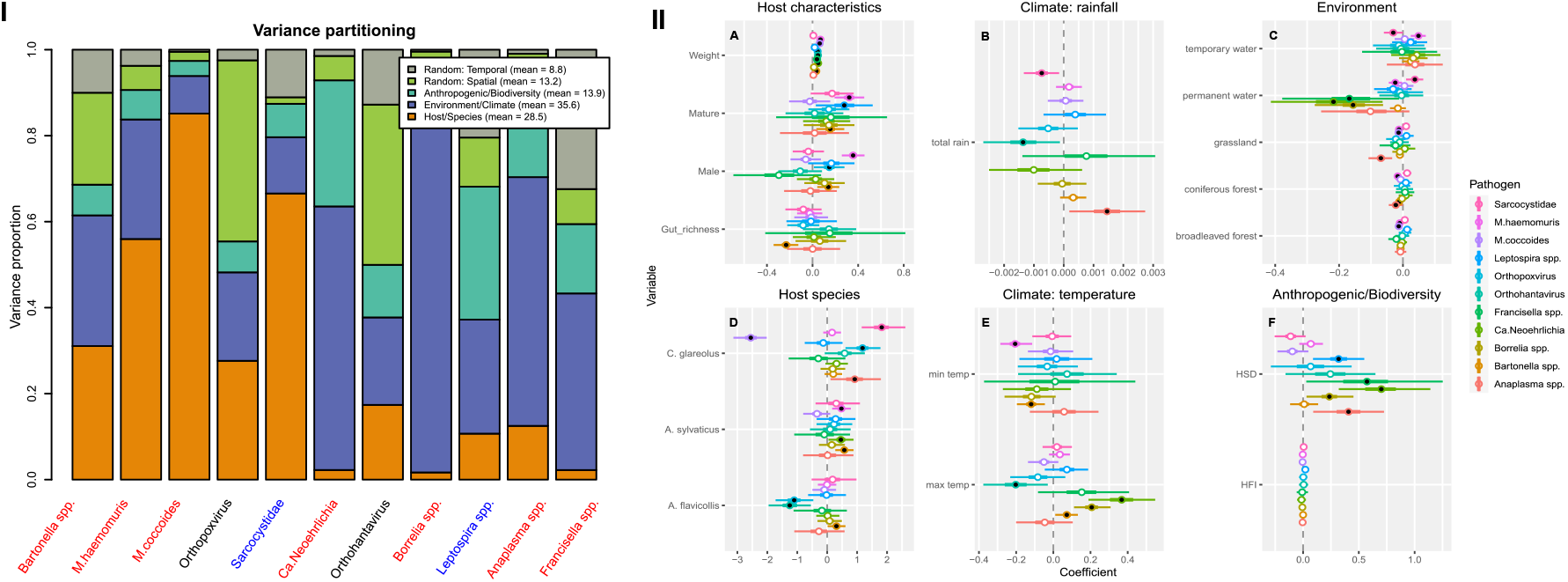
**I**. Variance partitioning for grouped fixed and random effects proportional to one from the JSDM INT+EXT+ANT. Pathogens have been ordered by decreasing values of variance explained and color coded with respect to main transmission mode (red = vector, black = direct and blue = environmental) **II**. Fixed effect estimates from JSDM model INT+EXT+ANT for host intrinsic factors [A, D], extrinsic factors such as different climatic conditions [B, E] and environmental conditions [C] and anthropogenic factors as the human footprint index (HFI) and host species diversity (HSD) (F). Lines indicate the 95% posterior probability and black dots indicate significant effects.

At the level of the linear predictor, the extrinsic environmental and climatic factors were found to be the primary drivers explaining the proportional variation in pathogen occurrences with 36% variance explained [Figure 6]. Host intrinsic factors followed with a total of 28%. Notably the vector-transmitted zoonotic pathogens (*Ca. Neoehrlichia, Borrelia* and *Anaplasma*) had between 59 – 85% of explained variance by climatic and environmental conditions. For *Francisella* and *Bartonella* spp. a similar, but less pronounced pattern was observed, with climatic and environmental conditions explaining 44% and 31% of proportional variance respectively. Anthropogenic variables were the most important drivers explaining observed variability in *Leptospira* spp. occurrences, while *M. haemomuris, M. coccoides* and *Sarcocystidae* were strongly dependent on host intrinsic characteristics. For the pathogens with a direct transmission mode, Orthohantavirus and Orthopoxvirus, the drivers were less pronounced. For Orthopoxvirus the random spatial latent variables accounted for a total of 43% of explained variance. For the Orthohantavirus the random spatial latent variable accounted for 38%, while the extrinsic factors contributed 19%, the intrinsic 17% and the anthropogenic 12%.

In general, a positive association between pathogen occurrence and heavier, more mature males was observed. *Bartonella* spp. was the sole pathogen showing a significant association with the gut-microbiome [Figure 6: II, A]. *A. flavicollis* was negatively associated with antiviral antibodies targeting Orthohantavirus and Orthopoxvirus, and positively with *Bartonella spp. A. sylvaticus* was positively associated with *Bartonella spp*., *Ca. Neoehrlichia* and *M. haemomuris*. The bacteria *M. coccoides* was virtually absent in *C. glareolus*, while an increased occurrence of *Sarcocystidae, Anaplasma* spp. and antiviral antibodies against Orthopoxvirus was associated with *C. glareolus* [Figure 6: II, D]. The percent coverage of grassland, forest and of permanent water bodies around a site was on average negatively associated with several pathogens, except those from the *Sarcosystidae*, which showed a positive tendency [Figure 6: II, C]. The occurrence of *Anaplasma* spp., *Bartonella* spp., and *M. haemomuris* was negatively associated with coniferous forests [Figure 6: II, C]. The minimum temperature during the past winter was negatively associated with *Bartonella* spp. and *M. haemomuris*, while the maximum temperature during the past summer showed opposing patterns for different pathogens but was mainly positively associated with vector transmitted pathogens [Figure 6: II, E]. Dryer conditions during the previous year seemed to favor directly transmitted Orthohanta- and Orthopoxviruses as well as *Sarcocystidae*, while wetter conditions favoured *Anaplasma spp*. [Figure 6: II, B]. The human footprint index was not significantly associated with pathogen occurrence, whereas an increase in host species diversity was associated with a significantly higher infection risk for five pathogens [Figure 6: II, F].

Trait-covariate association showed that directly transmitted pathogens associated negatively with the urban avoider *A. flavicollis*, while vector-based transmission was positively associated with individuals’ body weight. After accounting for covariates and transmission mode, residual associations remaining at the spatial level were predominantly positive, while a mix was observed at the temporal level (Supplemental Figure S2). At the spatial level positive residual association between vector transmitted *Bartonella* spp., *Anaplasma* spp., *Ca. Neoehrlichia, Mycoplasma* spp. and the environmentally transmitted *Leptospira* spp. were observed, while negative association between *Borrelia* spp., *Francisella* spp. and *Sarcocystidae* remained. Remaining residual co-occurrence at the temporal level showed both positive and negative residual associations indicating seasonal and annual differences need to be accounted for.

## Discussion

Small mammals are recognized as important reservoirs for zoonotic infections and known for their transmission potential to humans and livestock [12]. Here we present the largest European small mammal – zoonotic pathogen dataset empirically collected in a single study to date. We aimed to capitalize on this information to further improve our understanding of how different host intrinsic, extrinsic, and anthropogenic drivers impact host-pathogen communities and ultimately the risk of spillover.

### Host community

On average, host community specific richness was comparable between the two habitats, but variable between countries (Figure 2). This contrasts with the original expectation [54], but aligns with the study of Luza et al., 2021 [55], who found balanced species richness in human-modified habitats despite reduced functional diversity, notably because of local extinctions and immigration in western European temperate forest regions [55]. Host community turnover was associated with a gradient in anthropization as proportionally more urban dwellers were observed in more anthropized areas [Supplemental Figure S3]. Similarly, in South-East Asia it was shown that human-altered landscapes favour habitat generalists and synanthropic rodents [56]. It has been demonstrated that urbanization leads to a shift in species composition, with a selection towards those species that tend to have fast pace-of-life and are good dispersers [57]. In our dataset, no evidence was found for a shift towards faster pace-of-life species along the gradient, given the subset of small terrestrial mammals recorded, but differences in species composition were evident.

### Pathogen detection and co-infection/co-exposure

Small mammal host communities were exposed to a large variety of pathogens in both temperate European forests as well as urban green spaces. The three most prevalent pathogens with zoonotic potential belonged to two different genera: *Bartonella, Orthopoxvirus* spp., and to the family *Sarcocystidae*.

*Bartonella* spp. are reemerging ubiquitous bacteria that belong to a diverse group of Gram-negative, facultative intracellular pathogens known to cause endocarditis in humans [58]. Bartonellae inhabits both the gut of bloodsucking arthropod vectors including but not limited to fleas, lice, ticks, and sandflies as well as the bloodstream of mammalian hosts. Preliminary sequencing analysis of the gltA and rpoB genes on 11 positive individuals from France included in this study revealed the presence of six different *Bartonella* species including *Bartonella birtlesii, B. doshiae, B. gliris, B. taylorii*, one undetermined *Bartonella spp*. and *B. grahamii*, which is known to be zoonotic to humans.

Nearly 16% of specimens had detectable anti-Orthopoxvirus antibodies. The *Orthopoxvirus* genus includes species zoonotic diseases agents like smallpox, monkeypox and cowpox viruses, which are emergent since the cessation of smallpox vaccination. However, due to serological cross-reactivity it was not determined if the infections belonged specifically to those zoonotic agents. Infection occurred predominantly in *C. glareolus* and *R. norvegicus* but was also observed in the genus *Apodemus* and *Microtus* at lower prevalence. This confirms reports published earlier showing that Orthopoxvirus is endemic to Western Europe where bank voles, field voles and wood mice were described as important reservoir hosts [59], while *R. norvegicus* was shown to transmit cowpox virus to monkeys in the Netherlands [60].

*Sarcocystidae* spp. were detected predominantly in *C. glareolus* and observed at low prevalence in all genera, except in *M. musculus* and *Glis glis*. Blasting the 16s rRNA OTU showed 100% match with both *Neospora caninum* and its close relative *Toxoplasma gondii* as well as *Eimeria meleagrimitis. Neospora caninum* is not considered a zoonotic disease but can cause neosporosis in cattle with abortion as the prime clinical manifestation [61] and hind limb paralysis in dogs. Toxoplasmosis, with *Toxoplasma gondii* as its causative agent, usually results in absent or mild symptoms except in immunocompromised people and pregnant woman [62]. *Eimeria meleagrimitis* can cause mild disease in young turkeys [63].

Co-infection and co-exposure were commonplace, with up to seven pathogens detected simultaneously in one host during the present study. Positive non-random co-occurrences with *Bartonella* spp., which may be acting as symbionts in rodents [64], endoparasitic *Mycoplasma* spp. and several potentially zoonotic pathogens illustrate the importance of studying pathogen communities in a single host, as this can affect disease outcomes and transmissibility [65].

### Hosts-pathogen network

We showed that anthropization affects network properties. Anthropization is associated with increased fragmentation, urbanization and a shift in host community composition towards more synanthropic small mammals. Therefore we expected a more specialized network in the forested habitat, while more generalist host-pathogen interactions with the increasing gradient of urbanization were expected [23]. Our results were generally in line with these predictions as we observed a more specialized network in the temperate forest. Indeed, network-wide specialization (H2) is a metric that decreases as specialization increases, and was shown not to be affected by network sampling intensity [45]. A more connected network was observed in the urban green spaces with an increase in specialization asymmetry in accordance with more generalist host-pathogen interactions. Network structure was similar between the two habitats with four network modules each, but overall modularity was lower in urban green spaces. In south-east Asia, a study on rodent-helminth networks also found it to affect network properties, but not species richness of parasite communities per se [66]. Contrary to their findings where the gradient of fragmentation induced less connected and more modular rodent-helminth interactions, our study indicates an opposite pattern. Another study from the same area found rodent-tick networks to be more connected and showed lower modularity in an urban setting, whereas the pattern for rodent-microbes was opposite [19]. These differences highlight the complex suit of interaction at play in forming these networks. Differences in pathogen properties such as transmission traits and complexity of the life cycle, but also between temperate and tropical biomes regarding habitat fragmentation and host assemblages [15] may drive these observed variations.

Network analyses revealed a more specialized network with lower connectance in the temperate forest and important differences in key host species between habitats. In the urban green spaces, our analysis suggests the wood mouse (*Apodemus sylvaticus*) and the brown rat (*Rattus norvegicus*) as connector species, whereas the bank vole (*Clethrionomys glareolus*) and the yellow-necked mouse (*Apodemus flavicollis*) were identified in the temperate forests. This confirms our expectation that hosts with high centrality in pathogen networks in urban green spaces are urban dwellers or urban adapters, whereas in the temperate forests hosts with high centrality tend to avoid human settlements.

A depauperated assemblage of vector-transmitted zoonotic pathogens was observed in the urban host-pathogen network. Pathogens such as *Ehrlichia, Anaplasma, Orientia and Rickettsia* were absent which is in line with reports that synanthropic rodents such as *Rattus* and *Mus* species are seldom infested with ticks [67,68], and preferred hosts for reproduction such as ungulates are missing from urban environments. This highlights the potential impact of anthropization on the composition and diversity of zoonotic pathogens, with certain species being excluded or having a limited presence in urban environments. A similar pattern was observed in South-East Asia where tick infestation on *R. rattus* decreased along a rural-urban gradient. However, environmentally transmitted diseases such as *Leptospira* spp., and *T. gondii* showed elevated risks in habitats where synanthropic species thrive [19]. Urban encroachment into peri-urban wilderness and the increased presence of invasive species are likely to increase the suite of relevant host species. Understanding the absence or reduced occurrence of specific pathogens in urban areas is crucial for assessing disease risks and implementing effective control measures, as well as an in-depth assessment of the relative importance of abiotic factors and host species composition that drive patterns in pathogen occurrence.

### Pathogens community structure & drivers

We examined how different drivers shape pathogen communities in terrestrial small mammals along a gradient of anthropization. We focused on the most abundant host genera across the gradient, namely *Apodemus* spp. and *Clethrionomys glareolus*, as they occurred in all countries. Host species and host intrinsic factors explained most of the proportional variance in individual parasite occurrences for the bacterial *Mycoplasmas* and the protozoan *Sarcocystidae*. This is in line with several studies demonstrating the importance of individual host characteristics such as sex, maturity, and weight in parasite load [69] and parasite-host specificity on host-pathogen community assembly [70]. Notably, *Bartonella* was the only pathogen where occurrence correlated (negatively) with gut microbiome richness, following the hypothesis that an increase in gut microbiome richness is associated with increased host fitness and potentially resulting in a decrease in pathogen infection [71,72]. Our results also confirmed the critical role of extrinsic environmental and climatic factors in shaping pathogen community structure. This was especially detected for the vector-borne diseases, perhaps reflecting the strong impact of abiotic features, in particular temperature, dryness and land cover, on ectoparasite distribution and host-parasite interactions [73].

In line with the host-diversity-begets-parasite diversity relationship [8], our final model confirmed that pathogen communities get more diverse with an increase in overall rodent and vole biodiversity. On the other hand, anthropization (assessed via the human footprint index) was not associated with increased infection risk in our data. This suggested that the declining host diversity and consequent shift in host community competence usually associated with urbanization would not lead to an increase in disease burden in low-diversity habitats. Similarly, in a worldwide study on human infectious diseases, the disease burden was not found to be correlated with levels of biodiversity, consequently the effect of changing biodiversity on public health remains to be demonstrated [74]. Future studies should aim to integrate human case data, following a one-health approach, which would be a necessary next step to identify the factors associated with human-wildlife disease transmission.

#### Study limitations

Whereas we present the largest single-project study on rodents and rodent-borne diseases to date, based on standardized trapping protocol for terrestrial small mammals, bias in sampled rodent species was inevitable. Due to different ethical requirements, shrew species for example could not be trapped in all locations. Notably *Rattus norvegicus*, was either trapped with a larger single-case trap or with the help of pest control managers, but this was not performed in all countries or at an equal level. Therefore, in Ireland and Germany those species were absent from the urban area samples, which introduces biases. We have alleviated this in the HMSC model by restricting our analysis to the genera *Apodemus* and *Clethrionomys*, and by using an existing host species distribution model for diversity instead of relying on our own collections. In addition, as our protocol was designed with pathogen occurrence in mind, we could not estimate the densities of different host species.

Pathogen detection was also hampered by several limitations. In particular, the taxonomic resolution provided by the different approaches implemented may vary from strain (e.g. *Mycoplasma* spp. using 16S metabarcoding) to genus (e.g. IFA due to cross-reactivity) or even family (e.g. *Sarcocystidae* using 16S metabarcoding). The pathogen taxa included in our analyses may therefore not be specifically determined or may correspond to a diversity of species (e.g. coinfection of *Bartonella* species in the same host).

## Conclusion

We showed that a suite of complex biotic and abiotic interactions shape host-pathogen communities throughout Europe. Contrary to current emphasis on the relationship between biodiversity loss and dilution we demonstrate that host intrinsic characteristics, local habitat and climatic filtering are key factors that drive pathogen communities at the local landscape level. Anthropization did not affect pathogen occurrences but affected host-pathogen network properties. By comprehensively studying these filtering processes, we can gain a better understanding of the drivers behind pathogen variations, ultimately informing proactive measures to mitigate zoonotic disease risks and safeguard public health.

## Data availability statement

Raw sequencing data for the 16Sv4 rRNA gene from spleen and colon of small mammal samples are deposited in ZENODO, alongside scripts for analyses [respectively https://doi.org/10.5281/zenodo.12518286 and https://doi.org/10.5281/zenodo.12527197].

Geographical data, covariates and information on individual specimen used for the analyses in the paper, are deposited alongside scripts for analyses and figures on github [https://github.com/vsluydts/bioroddis].

## Acknowledgments

We thank all assistants, technicians and students for their valuable help in the field and laboratory work. We thank Annetta Zintl for the valuable discussion about *Anaplasma* spp.

## Author contributions

**Vincent Sluydts**: C, DC, FA, I, M, R, VA, WO; **Marie Bouilloud**: FA, M, R, WR. **Maxime Galan**: C, DC, FA, M, R, I, SU, WR; **Hussein Alburkat**: R, WR; **Anaïs Bordes**: R; **Vincent Bourret**: R, WR; **Valeria Colombo**: R, WR; **Luc DeBruyn**: M, R, WR; **Lara Dutra**: R, WR; **Jana Eccard**: FU, SU, R, WR; **Jasmin Firozpoor**: R, WR; **Romain Gallet**: R, WR, **Maciej Grzybek**: FU, R, WR; **Heikki Henttonen**: C, WR; **Jens Jacob**: R, WR; **Andrew McManus**: R, WR ; **Tarja Sironen**: FU, R, SU, WR; **Peter Stuart:** FU, R, SU, WR; **Caroline Tatard**: R, WR; **Benjamin Roche**: FU, R, C, M, SU, WR; **Herwig Leirs**: FU, R, SU, WR**; Nathalie Charbonnel**: FU, C, M, PA, SU, WR. Abbreviations follow the Elsevier author credit classification.

## References

1. Keesing, F. and Ostfeld, R.S. (2021) Impacts of biodiversity and biodiversity loss on zoonotic diseases. Proc Natl Acad Sci U S A 118. 10.1073/pnas.2023540118

2. Johnson, P.T. et al. (2015) Frontiers in research on biodiversity and disease. Ecol Lett 18, 1119–1133. 10.1111/ele.12479

3. Estrada-Pena, A. et al. (2014) Effects of environmental change on zoonotic disease risk: an ecological primer. Trends Parasitol 30, 205–214. 10.1016/j.pt.2014.02.003

4. Rynkiewicz, E.C. et al. (2019) Linking community assembly and structure across scales in a wild mouse parasite community. Ecol Evol 9, 13752–13763. 10.1002/ece3.5785

5. Leibold, M.A. et al. (2021) The internal structure of metacommunities. Oikos 2022. 10.1111/oik.08618

6. Leibold, M.A. and Chase, J.M. (2018) Metacommunity Ecology (Vol. 59, Princeton University Press (Monograph)

7. Johnson, P.T. et al. (2015) Why infectious disease research needs community ecology. Science 349, 1259504. 10.1126/science.1259504

8. Johnson, P.T. et al. (2016) Habitat heterogeneity drives the host-diversity-begets-parasite-diversity relationship: evidence from experimental and field studies. Ecol Lett 19, 752–761. 10.1111/ele.12609

9. Halliday, F.W. et al. (2020) Biodiversity loss underlies the dilution effect of biodiversity. Ecol Lett 23, 1611–1622. 10.1111/ele.13590

10. Suzán, G. et al. (2015) Metacommunity and phyologenetic structure determine wildlife and zoonotic infectious disease patterns in time and space. Ecology and Evolution 5, 865–873. 10.1002/ece3.1404

11. Watson, J.E.M. and Venter, O. (2019) Mapping the continuum of humanity’s footprint on land. One Earth 1, 175–180. 10.1016/j.oneear.2019.09.004

12. Han, B.A. et al. (2015) Rodent reservoirs of future zoonotic diseases. Proc Natl Acad Sci U S A 112, 7039–7044. 10.1073/pnas.1501598112

13. Bradley, C.A. and Altizer, S. (2007) Urbanization and the ecology of wildlife diseases. Trends Ecol Evol 22, 95–102. 10.1016/j.tree.2006.11.001

14. Jones, K.E. et al. (2008) Global trends in emerging infectious diseases. Nature 451, 990–993. 10.1038/nature06536

15. Gibb, R. et al. (2020) Zoonotic host diversity increases in human-dominated ecosystems. Nature 584, 398–402. 10.1038/s41586-020-2562-8

16. Murray, M.H. et al. (2019) City sicker? A meta‐analysis of wildlife health and urbanization. Front Ecol Env 17, 575–583. 10.1002/fee.2126

17. Albery, G.F. et al. (2022) Urban-adapted mammal species have more known pathogens. Nat Ecol Evol 6, 794–801. 10.1038/s41559-022-01723-0

18. Methot, P.O. and Alizon, S. (2014) What is a pathogen? Toward a process view of host-parasite interactions. Virulence 5, 775–785. 10.4161/21505594.2014.960726

19. Blasdell, K.R. et al. (2022) Rats and the city: Implications of urbanization on zoonotic disease risk in Southeast Asia. Proc Natl Acad Sci U S A 119, e2112341119. 10.1073/pnas.2112341119

20. Himsworth, C.G. et al. (2013) Rats, cities, people, and pathogens: a systematic review and narrative synthesis of literature regarding the ecology of rat-associated zoonoses in urban centers. Vect Borne Zoon Dis 13, 349–359. 10.1089/vbz.2012.1195

21. Poulin, R. (2010) Network analysis shining light on parasite ecology and diversity. Trends Parasitol 26, 492–498. 10.1016/j.pt.2010.05.008

22. Runghen, R. et al. (2021) Network analysis: Ten years shining light on host-parasite interactions. Trends Parasitol 37, 445–455. 10.1016/j.pt.2021.01.005

23. Morand, S. et al. (2019) Changing landscapes of Southeast Asia and rodent‐borne diseases: decreased diversity but increased transmission risk. Ecol Appl 29 (4). 10.1002/eap.1886

24. Urbieta, G.L. et al. (2021) Modularity and specialization in bat-fly interaction networks are remarkably consistent across patches within urbanized landscapes and spatial scales. Curr Zool 67, 403–410. 10.1093/cz/zoaa072

25. Pradel, J. et al. (2022) Small terrestrial mammals (Rodentia and Soricomorpha) along a gradient of forest anthropisation (reserves, managed forests, urban parks) in France. Biodivers Data J 10, e95214. 10.3897/BDJ.10.e95214

26. Bayerns, S.N.S. (2024). Rodent composition of urban and forested areas in Potsdam, Germany.

27. Kallio-Kokko, H. et al. (2006) Hantavirus and arenavirus antibody prevalence in rodents and humans in Trentino, Northern Italy. Epidemiol Infect 134, 830–836. 10.1017/S0950268805005431

28. Dobigny, G. et al. (2015) Urban market gardening and rodent-borne pathogenic leptospira in arid zones: A case study in Niamey, Niger. PLoS Negl Trop Dis 9, e0004097. 10.1371/journal.pntd.0004097

29. Kozich, J.J. et al. (2013) Development of a dual-index sequencing strategy and curation pipeline for analyzing amplicon sequence data on the MiSeq Illumina sequencing platform. Appl Environ Microbiol 79, 5112–5120. 10.1128/AEM.01043-13

30. Galan, M. et al. (2016) 16S rRNA amplicon sequencing for epidemiological surveys of bacteria in wildlife. mSystems 1. 10.1128/mSystems.00032-16

31. Escudie, F. et al. (2018) FROGS: Find, Rapidly, OTUs with Galaxy Solution. Bioinformatics 34, 1287–1294. 10.1093/bioinformatics/btx791

32. Wang, Q. et al. (2007) Naive Bayesian classifier for rapid assignment of rRNA sequences into the new bacterial taxonomy. Appl Environ Microbiol 73, 5261–5267. 10.1128/AEM.00062-07

33. Camacho, C. et al. (2009) BLAST+: architecture and applications. BMC Bioinformatics 10, 421. 10.1186/1471-2105-10-421

34. Abbate, J.L. et al. (2024) Pathogen community composition and co-infection patterns in a wild community of rodents Peer Community Journal 4 (e14). 10.24072/pci.ecology.100087

35. Reese, A.T. and Dunn, R.R. (2018) Drivers of Microbiome Biodiversity: A Review of General Rules, Feces, and Ignorance. mBio 9. 10.1128/mBio.01294-18

36. Bouilloud, M. (2023) Three-way relationships between gut microbiota, helminth assemblages and bacterial infections in wild rodent populations. Peer Community Journal 3 (e18). 10.24072/pci.infections.106000

37. Vanwambeke, S.O. et al. (2019) Spatial dynamics of a zoonotic orthohantavirus disease through heterogenous data on rodents, rodent infections, and human disease. Sci Rep 9, 2329. 10.1038/s41598-019-38802-5

38. Sanderson, E.W. et al. (2022). The march of the human footprint. ecoevorxiv

39. Wint, G.R.W. (2013) Four Rodent and Vole Biodiversity Models for Europe. Journal of Open Public Health Data 1. 10.5334/jophd.ac

40. Cao, Y. et al. (1997) Similarity measure bias in river benthic Aufwuchs community analysis. Water Environment Research 69, 95–106. 10.2175/106143097×125227

41. Oksanen J et al. (2022). vegan: Community Ecology Package. R package version 2.6-4 ed.

42. Plourde, B.T. et al. (2017) Are disease reservoirs special? Taxonomic and life history characteristics. PLoS One 12, e0180716. 10.1371/journal.pone.0180716

43. Veech, J.A. and Peres-Neto, P. (2013) A probabilistic model for analysing species co-occurrence. Global Ecology and Biogeography 22, 252–260. 10.1111/j.1466-8238.2012.00789.x

44. Dormann, C.F. et al. (2008). Introducing the bipartite package: Analysing ecological networks. R News

45. Bluthgen, N. et al. (2006) Measuring specialization in species interaction networks. BMC Ecol 6, 9. 10.1186/1472-6785-6-9

46. Bluthgen, N. et al. (2008) What do interaction network metrics tell us about specialization and biological traits? Ecology 89, 3387–3399. 10.1890/07-2121.1

47. Patefield, W.M. (1981) An efficient method of generating random R x C tables with given row and column totals. Journal of the Royal Statistical Society. Series C (Applied Statistics) 30, 91–97

48. Martín González, A.M. et al. (2010) Centrality measures and the importance of generalist species in pollination networks. Ecological Complexity 7, 36–43. 10.1016/j.ecocom.2009.03.008

49. Warton, D.I. et al. (2015) So Many Variables: Joint Modeling in Community Ecology. Trends Ecol Evol 30, 766–779. 10.1016/j.tree.2015.09.007

50. Ovaskainen, O. et al. (2017) How to make more out of community data? A conceptual framework and its implementation as models and software. Ecol Lett 20, 561–576. 10.1111/ele.12757

51. Tjur, T. (2009) Coefficients of determination in logistic regression models—A new proposal: The coefficient of discrimination. The American Statistician 63, 366–372. 10.1198/tast.2009.08210

52. Tikhonov, G. et al. (2020) Joint species distribution modelling with the r-package Hmsc. Methods Ecol Evol 11, 442–447. 10.1111/2041-210X.13345

53. Fischer, J.D. et al. (2015) Categorizing wildlife responses to urbanization and conservation implications of terminology. Conserv Biol 29, 1246–1248. 10.1111/cobi.12451

54. Cavia, R. et al. (2009) Changes in rodent communities according to the landscape structure in an urban ecosystem. Landscape and Urban Planning 90, 11–19. 10.1016/j.landurbplan.2008.10.017

55. Luza, A.L. et al. (2021) Functional redundancy of non‐volant small mammals increases in human‐modified habitats. J Biogeography 48, 2967–2980. 10.1111/jbi.14264

56. Morand, S. et al. (2019) Changing landscapes of Southeast Asia and rodent-borne diseases: decreased diversity but increased transmission risks. Ecological Applications 29. 10.1002/eap.1886

57. Santini, L. et al. (2019) One strategy does not fit all: determinants of urban adaptation in mammals. Ecol Lett 22, 365–376. 10.1111/ele.13199

58. Okaro, U. et al. (2017) Bartonella Species, an Emerging Cause of Blood-Culture-Negative Endocarditis. Clin Microbiol Rev 30, 709–746. 10.1128/CMR.00013-17

59. Kinnunen, P.M. et al. (2011) Orthopox virus infections in Eurasian wild rodents. Vector Borne Zoonotic Dis 11, 1133–1140. 10.1089/vbz.2010.0170

60. Martina, B.E.E. et al. (2006) Cowpox virus transmission from rats to monkeys, the Netherlands. Emerg Infect Dis 12, 1005–1007. 10.3201/eid1206.051513

61. Donahoe, S.L. et al. (2015) A review of neosporosis and pathologic findings of Neospora caninum infection in wildlife. Int J Parasitol Parasites Wildl 4, 216–238. 10.1016/j.ijppaw.2015.04.002

62. Ybanez, R.H.D. et al. (2020) Review on the Current Trends of Toxoplasmosis Serodiagnosis in Humans. Front Cell Infect Microbiol 10, 204. 10.3389/fcimb.2020.00204

63. Gadde, U.D. et al. (2020) Pathology caused by three species of Eimeria that infect the turkey with a description of a scoring system for intestinal lesions. Avian Pathol 49, 80–86. 10.1080/03079457.2019.1669767

64. McKee, C.D. et al. (2021) Bats are key hosts in the radiation of mammal-associated Bartonella bacteria. Infect Genet Evol 89, 104719. 10.1016/j.meegid.2021.104719

65. Hoarau, A.O.G. et al. (2020) Coinfections in wildlife: Focus on a neglected aspect of infectious disease epidemiology. PLoS Pathog 16, e1008790. 10.1371/journal.ppat.1008790

66. Bordes, F. et al. (2015) Habitat fragmentation alters the properties of a host-parasite network: rodents and their helminths in South-East Asia. J Anim Ecol 84, 1253–1263. 10.1111/1365-2656.12368

67. Hornok, S. et al. (2015) Synanthropic rodents and their ectoparasites as carriers of a novel haemoplasma and vector-borne, zoonotic pathogens indoors. Parasit Vectors 8, 27. 10.1186/s13071-014-0630-3

68. Mihalca, A.D. et al. (2012) Tick parasites of rodents in Romania: host preferences, community structure and geographical distribution. Parasites & Vectors 5, 1–7. 10.1186/1756-3305-5-266

69. Hillegass, M.A. et al. (2008) The influence of sex and sociality on parasite loads in an African ground squirrel. Behavioral Ecol 19, 1006–1011. 10.1093/beheco/arn070

70. Dallas, T. and Presley, S.J. (2014) Relative importance of host environment, transmission potential and host phylogeny to the structure of parasite metacommunities. Oikos 123, 866–874. 10.1111/oik.00707

71. Suzuki, T.A. (2017) Links between Natural Variation in the Microbiome and Host Fitness in Wild Mammals. Integr Comp Biol 57, 756–769. 10.1093/icb/icx104

72. Zaneveld, J.R. et al. (2017) Stress and stability: applying the Anna Karenina principle to animal microbiomes. Nat Microbiol 2, 17121. 10.1038/nmicrobiol.2017.121

73. Poisot, T. et al. (2017) Hosts, parasites and their interactions respond to different climatic variables. Global Ecol Biogeography 26, 942–951. 10.1111/geb.12602

74. Wood, C.L. et al. (2017) Human infectious disease burdens decrease with urbanization but not with biodiversity. Philos Trans R Soc Lond B Biol Sci 372. 10.1098/rstb.2016.0122

