## Supplemental table and figures for "Drivers of host-pathogen community assemblies in European forests and urban green spaces"

**APPENDIX**

**Supplementary Table S1.** Pathogens observed in the small mammal host communities ordered by Phylum and prevalence. * indicates genera with zoonotic potential.

| Phylum | Disease | Genus | Transmission | Infected (n) | Prevalence (%) | Detection method (sample) |
|---|---|---|---|---|---|---|
| Bacteria | Endocarditits* | Bartonella | Vector | 1812 | 46.94 | 16S - Spleen |
| Bacteria | Rodent disease | Mycoplasma haemomuris | Vector | 1460 | 37.82 | 16S - Spleen |
| Bacteria | Rodent disease | Mycoplasma coccoides | Vector | 679 | 17.59 | 16S - Spleen |
| Bacteria | Ehrlichiosis* | Ehrlichia | Vector | 368 | 9.53 | 16S - Spleen |
| Bacteria | Lyme disease* | Borrelia | Vector | 134 | 3.47 | 16S - Spleen |
| Bacteria | Leptospirosis* | Leptospira | Indirect (water) | 128 | 3.32 | 16S + PCR - Spleen/Kidney |
| Bacteria | Anaplasmosis* | Anaplasma | Vector | 77 | 1.99 | 16S - Spleen |
| Bacteria | Tularemia* | Francisella | Vector | 69 | 1.79 | 16S - Spleen |
| Bacteria | Scrub typhus* | Orientia | Vector | 29 | 0.75 | 16S - Spleen |
| Bacteria | Rodent disease | Mycoplasma insons | Vector | 25 | 0.65 | 16S - Spleen |
| Bacteria | Rodent disease | Mycoplasma ravipulmonis | Vector | 21 | 0.54 | 16S - Spleen |
| Bacteria | Rodent disease* | Mycoplasma penetrans | Vector | 21 | 0.54 | 16S - Spleen |
| Bacteria | Chlamydia* | Chlamydia | Direct | 18 | 0.47 | 16S - Spleen |
| Bacteria | Rat-bite fever* | Streptobacillus | Direct | 11 | 0.28 | 16S - Spleen |
| Bacteria | Spotted fever, Thyphus* | Rickettsia | Vector | 9 | 0.23 | 16S - Spleen |
| Bacteria | Rodent disease | Mycoplasma microti | Vector | 6 | 0.16 | 16S - Spleen |
| Protozoa | Sarcocystosis* | Sarcocystidae | Indirect (Sporocysts) | 395 | 10.23 | 16S - Spleen |
| Viruses | Cowpox, Smallpox, Monkeypox* | Orthopoxvirus | Direct (bites) | 610 | 15.8 | IFA - Blood |
| Viruses | Hemorrhagic Fever with Renal Syndrome* | Orthohantavirus | Direct (air, bites) | 189 | 4.9 | IFA - Blood |
| Viruses | Lymphocytic Choriomeningitis (LCMV)* | Mammarenavirus | Direct (air, bites) | 35 | 0.91 | IFA - Blood |

**Supplementary Figure S1.** Co-occurrences as co-infection or co-exposure with indication of pairs of infection observed more (+) or less than expected ad random.

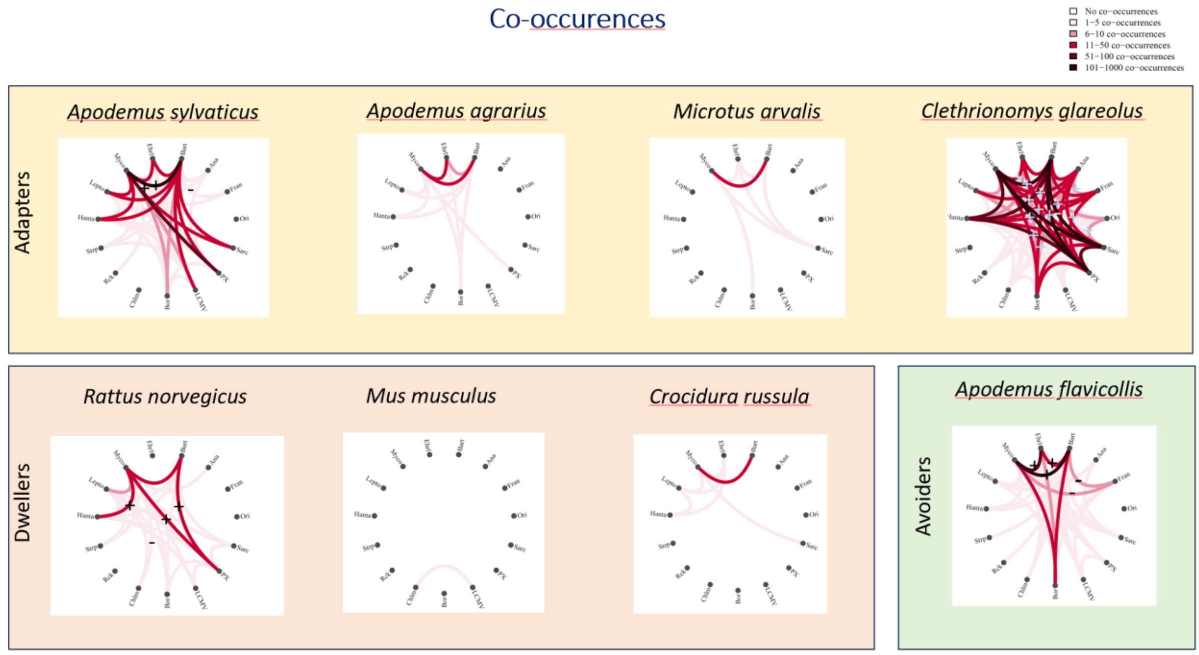

**Supplementary Figure S2.** A. Trait-covariate association at a 95% posterior probability level from the JSDM model with intrinsic, extrinsic, and anthropogenic factors included [INT + EXT+ ANT]. B. Residual random associations from the same model at the spatial and C. temporal level.

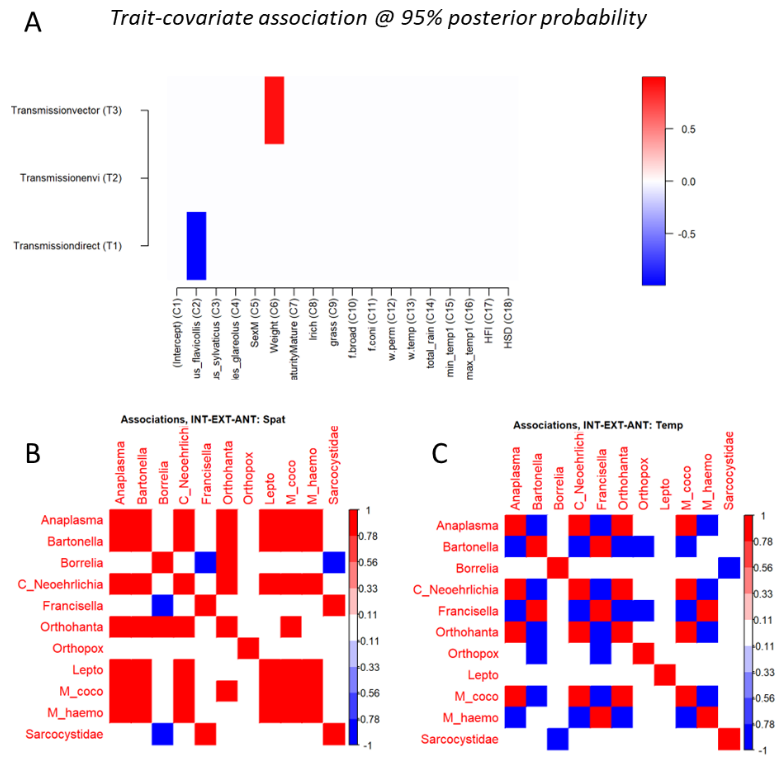


**Supplementary Figure S3**: Anthropization gradient versus host species categorisation in urban dweller, avoider adapter.

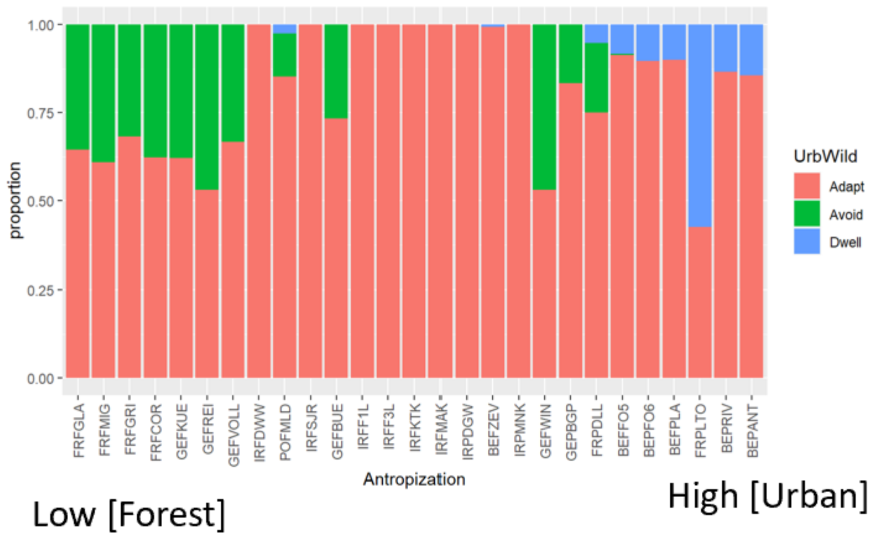
